# Determining significant correlation between pairs of extant characters in a small parsimony framework

**DOI:** 10.1101/2021.01.26.428213

**Authors:** Kaustubh Khandai, Cristian Navarro-Martinez, Brendan Smith, Rebecca Buonopane, S. Ashley Byun, Murray Patterson

**Author notes:** Joint authors (S. Ashley Byun, Murray Patterson).

## Abstract

When studying the evolutionary relationships among a set of species, the principle of parsimony states that a relationship involving the fewest number of evolutionary events is likely the correct one. Due to its simplicity, this principle was formalized in the context of computational evolutionary biology decades ago by, *e.g.*, Fitch and Sankoff. Because the parsimony framework does not require a model of evolution, unlike maximum likelihood or Bayesian approaches, it is often a good starting point when no reasonable estimate of such a model is available.

In this work, we devise a method for determining if pairs of discrete characters are significantly correlated across all most parsimonious reconstructions, given a set of species on these characters, and an evolutionary tree. The first step of this method is to use Sankoff’s algorithm to compute *all* most parsimonious assignments of ancestral states (of each character) to the internal nodes of the phylogeny. Correlation between a pair of evolutionary events (*e.g.*, absent to present) for a pair of characters is then determined by the (co-) occurrence patterns between the sets of their respective ancestral assignments. The probability of obtaining a correlation this extreme (or more) under a null hypothesis where the events happen randomly on the evolutionary tree is then used to assess the significance of this correlation. We implement this method: `parcours` (PARsimonious CO-occURrenceS) and use it to identify significantly correlated evolution among vocalizations and morphological characters in the Felidae family.

The `parcours` tool is freely available at https://github.com/murraypatterson/parcours

## 1 Introduction

The principle of parsimony first appeared in the context of computational evolutionary biology in [12, 4, 15, 20]. Following this, Sankoff and Rousseau generalized this to allow the association of a (different) cost to each transition between a pair of states [40, 41]. A few years after that, Felsenstein noticed that parsimony could produce misleading results when the evolutionary rate of change on the branches of the phylogeny is high [16]. Because parsimony does not take branch length into account, it ignores the fact that many changes on a long branch — while being far from parsimonious — may not be so unlikely in this case, resulting in a bias coined as “long branch attraction”. This paved the way for a proposed refinement to parsimony known as the maximum likelihood method [16, 17]. Because of an explosion in molecular sequencing data, and the sophisticated understanding of the evolutionary rate of change in this setting, maximum likelihood has become the de facto framework for inferring phylogeny. Many popular software tools which implement maximum likelihood include RAxML [43], PHYLIP [18] and the Bio++ library [21]. Even more recently, some Bayesian approaches have appeared, which sample the space of likely trees using the Markov-Chain Monte Carlo (MCMC) method, resulting in tools such as MrBayes [23] and Beast [10].

However, in cases where there is no model of evolution, or no reasonable estimation of the rate of evolutionary change on the branches of the phylogeny, parsimony is a good starting point. In fact, even in the presence of such a model, there are still conditions under which a maximum likelihood phylogeny is always a maximum parsimony phylogeny [47, 45]. Finally, when the evolutionary rate of change of different characters is heterogeneous, *i.e.*, there are (different) rates of change, maximum parsimony has even been shown to perform *substantially better* than maximum likelihood and Bayesian approaches [27]. A good example is cancer phylogenetics [22, 42, 14, 7], where little is known about the modes of evolution of cancer cells in a tumor. Due to its importance, this setting has seen a renewed interest in high-performance methods for computing maximum parsimonies in practice, such as SCITE [25] and SASC [6].

The setting of the present work is an evolutionary study of vocalizations and morphological characters [5] among members of the family Felidae. Felids possess a range of intraspecific vocalizations for close, medium, and long range communication. These vocalizations can vary from discrete calls such as the spit or hiss to more graded calls such as the mew, main call, growl and snarl [46]. There is no evidence that felid vocalizations are learned: it is more likely that these calls are genetically determined [35, 13, 38]. While there are 14 major discrete and graded calls documented in Felidae, not all calls are produced by all species [36]. In [37], the authors map some of these calls to a molecular phylogeny of Felidae, to show that it is consistent with what was previously known, strengthening the argument that vocalizations are genetically determined. Here we consider a similar approach, but because (a) an obvious model of evolution is lacking in this case; (b) the possibility that vocalizations within a given group of species can evolve at considerably different rates [37]; and (c) that rates for specific characters can differ between different lineages within that group [37], it follows from [27] that parsimony is more appropriate than maximum likelihood or a Bayesian approaches.

In this work, we develop a general framework to determine, among pairs of characters in a phylogeny, which of them are significantly correlated (or co-occurring), when no model of evolution is present. We then use this framework to understand how these vocalizations and morphological characters may have evolved within Felidae, how they might correlate (or have co-evolved) with each other, and which ones among these are significantly correlated. The first step of this approach is to infer, for each character, the set of *all* most parsimonious assignments of ancestral states (small parsimonies) in the phylogeny. Then, for each character, we construct from its set of small parsimonies, a *consensus map* (see Section 3) for each (type of) evolutionary event (*e.g.*, absent to present) along branches of the phylogeny. Correlation between a pair of evolutionary events (for a pair of characters) is then determined by how much their respective consensus maps overlap. Finally, this correlation is deemed significant if it is higher than expected in a null model where the evolutionary events happen randomly on the tree. We implement this approach in a tool called `parcours`, and use it to detect significantly correlated evolution among felid vocalizations and morphological characters, obtaining results that are consistent with the literature [37] as well as some interesting associations. While various methods for detecting correlated evolution exist, they tend to use only *phylogenetic profiles* [30], or are based on maximum likelihood [2, 9, 33], where a model of evolution is needed. Methods that determine co-evolution in the parsimony framework exist as well [34, 11], however they are aimed at reconstructing ancestral gene *adjacencies*, given extant co-localization information. Finally, while some of the maximum likelihood software tools have a “parsimony mode”, *e.g.*, [18, 21], the character information must be encoded using restrictive alphabets, and there is no automatic way to compute *all* most parsimonious ancestral states for a character — something which is central to our framework. On the contrary, `parcours` takes character information as a *column-separated values* (CSV) file, inferring the alphabet from this input, and efficiently computes all small parsimonies. In summary, our contribution is a methodology for inferring correlation among pairs of characters when no model of evolution is available, assessing the significance of these associations, and implementing it in an open-source software that is easy enough to be used as a pedagogical tool.

This paper is structured as follows. In Sec. 2, we provide the background on parsimony, as well as our approach for efficiently computing *all* most parsimonious ancestral states in a phylogeny, given a set of extant states. In Sec. 3, we present our approach for computing correlation between pairs of characters from all such parsimonies. In Sec. 4, we describe how to compute the probability of obtaining a correlation as high (or higher) by chance. In Sec. 5, we describe an experimental analysis of the implementation, `parcours`, of our method on felid vocalizations and morphological characters, and then discuss the results in Sec 6. Finally, we conclude the paper in Sec. 7 with a discussion of future directions.

## 2 Small Parsimony

In computing a parsimony, the input is typically *character-based*, involving a set of species, each over a set of *characters* (*e.g.*, weight category), where each character can be in a number of *states* (*e.g.*, low, medium, high, unknown, etc.). The idea is that each species is in one of these states for each character — *e.g.*, in the Puma, the weight (a morphological character [5]) is “high” — and we want to understand what states the ancestors of these species could be in. Given a *phylogeny* on these species: a tree where all of these species are leaves, a *small parsimony* is an assignment of states to the internal (ancestral) nodes of this tree which minimizes the number of changes of state among the characters along branches of the tree.^1^ We illustrate this with the following example.

Suppose we have the four species: Puma, Jaguarundi, Cheetah (of the Puma lineage), and Pallas cat (outgroup species from the nearby Leopard cat lineage), alongside the phylogeny depicted in Fig. 1a, implying the existence of the ancestors *X*, *Y* and *Z*. We are given some character, *e.g.*, weight, which can be in one of a number of states taken from *alphabet* ∑ = {low, high, unknown}. If weight is high only in Puma and Cheetah as in Fig. 1b, then the assignment of low to all ancestors would be a small parsimony. This parsimony has two changes of state: a change low→high on the branches *Z*→Puma and *Y*→Cheetah — implying convergent increase of weight in these species. Another small parsimony is that weight is high in *Y* and *Z* (low in *X*) — implying that high weight was ancestral (in *Y*) to the Puma, Jaguarundi and Cheetah, and later decreased in the Jaguarundi. A principled way to infer all small parsimonies is to use Sankoff’s algorithm [40], which also makes use of a *cost matrix δ*, depicted in Fig. 1c, that encodes the evolutionary cost of each state change along a branch. Since the change low→high costs *δ*_low,high_ = 1, and vice versa (*δ*_high,low_ = 1), it follows that each of the small parsimonies mentioned above have an overall cost, or *score* of 2 in this framework. A simple inspection of all possibilities shows that 2 is the minimum score of any assignment of ancestral states.

**Figure 1:**
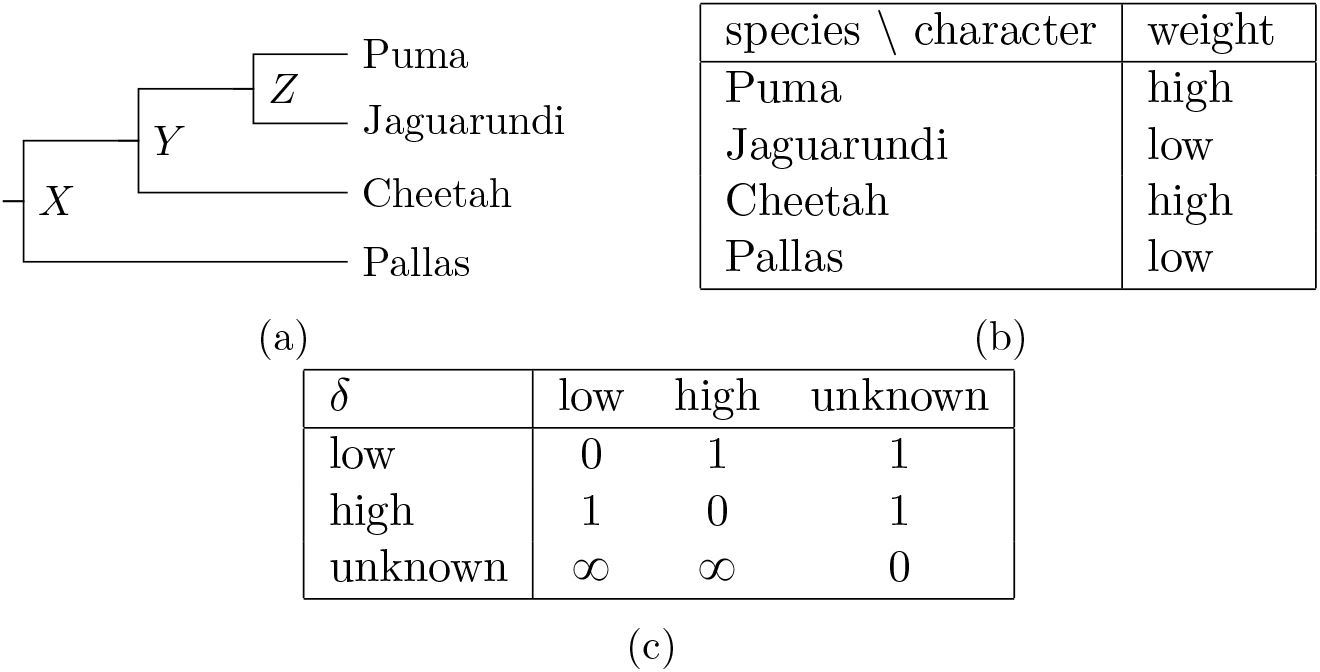
A (a) phylogeny and (b) the extant state of character weight in four species; and (c) the cost *δ_i,j_* of the change *i*→*j* from state *i* in the parent to state j in the child along a branch of the phylogeny, *e.g., δ*_high,unknown_ = 1 (*δ*_unknown,high_ = ∞).

Sankoff’s algorithm [40] computes all (small) parsimonies given a phylogeny and the extant states of a character in a set of species, and a cost matrix *δ*, *e.g.*, Fig. 1. The algorithm has a bottom-up phase (from the leaves to the root of the phylogeny), and then a top-down phase (from the root to the leaves). The bottom-up phase is to compute *s_i_*(*u*): the minimum score of an assignment of ancestral states in the subtree of the phylogeny rooted at node *u* when *u* has state *i* ∈ ∑, according to the recurrence:

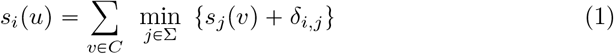

where *C* is the set of children of node *u* in the phylogeny. The idea is that the score is known for any extant species (a leaf node in the phylogeny), and is coded as *s_i_*(*u*) = 0 if species *u* has state *i*, and ∞ otherwise. The score for the internal nodes is then computed according to Eq. 1 in a bottom-up dynamic programming fashion, starting from the leaves. The boxes in Fig. 2 depict the result of this first phase on the instance of Fig. 1, for example,

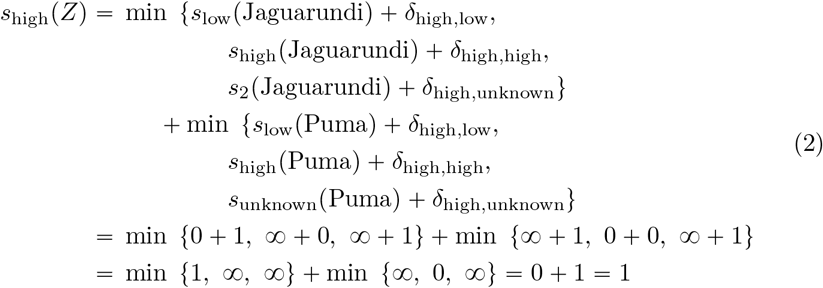

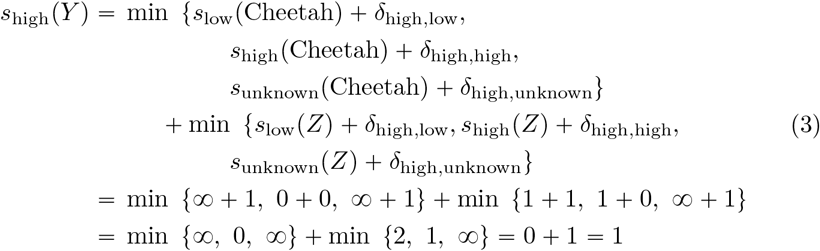

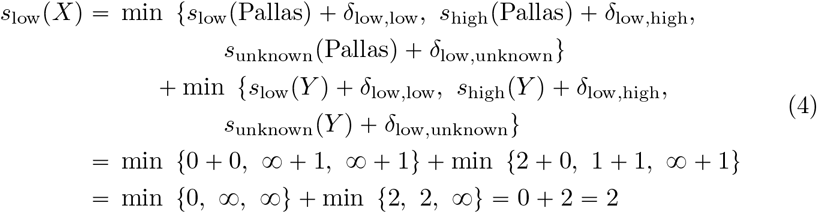

**Figure 2:**
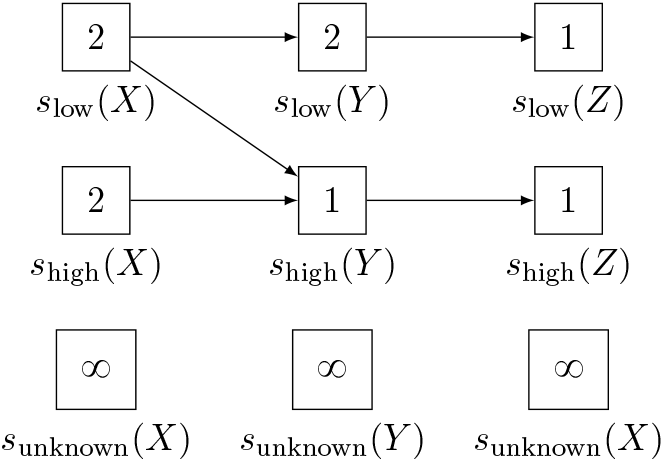
Graph structure resulting from Sankoff’s algorithm on the instance of Fig. 1. The *boxes* contain the values of *s_i_*(*u*) computed in the bottom-up phase, for each state *i* ∈ Σ and internal node *u* ∈ {*X, Y,Z*} of the phylogeny. The *arrows* connecting these boxes are computed in the top-down phase.

In general, for a given character with |Σ| = *k*, this procedure would take time *O*(*nk*), where *n* is the number of species, since we compute *nk* values, and computing each one takes (amortized) constant time.

After the bottom-up phase, we know the minimum score of any assignment of ancestral states (it is the minimum of those *s_i_* for *i* ∈ Σ, at the root), but we do not yet have an ancestral assignment of states. Here, since *X* is the root in Fig. 1a, we see from Fig. 2 that the minimum score for this example is 2, as we saw earlier. Note that this minimum may not be unique; indeed, here *s*_low_(*X*) = *s*_high_(*X*) = 2, meaning that in at least one parsimony, weight is low in *X*, and in at least one other parsimony, weight is high in *X* (but never unknown in any parsimony, *i.e., s*_unknown_(*X*) = ∞). Now, to reconstruct one of these ancestral assignments of minimum score — the top-down phase of Sankoff’s algorithm — we first assign to the root, one of the states *i* ∈ Σ for which *s_i_* is minimum. We then determine those states in each child of the root from which si can be derived (these may not be unique either), and assign those states accordingly. We continue, recursively, in a top-down fashion until we reach the leaves of the tree, having assigned all states at this point. For example, *s*_low_(*X*) = 2, and can be derived in Eq. 4 from *S*_high_(*Y*) (and *s*_low_(Pallas)), which is in turn derived in Eq. 3 from *s*_high_(*Z*) (and *s*_high_ (Cheetah)). This corresponds to the second parsimony mentioned earlier, where high weight was ancestral in *Y*, and later decreased in the Jaguarundi. Notice that *s*_low_(*X*) can also be derived in Eq. 4 from *s*_low_(*Y*), which is in turn derived from *s*_low_(*Z*). This is the first parsimony where weight is low in all ancestors.

One can compactly represent *all* parsimonies as a graph structure with a *box* for each si at each internal node of the phylogeny (*e.g.*, boxes of Fig. 2), and an *arrow* from box *s_i_*(*u*) to *s_j_* (*v*) for some node u and its child *v* in the phylogeny, whenever *s_i_*(*u*) can be derived from *s_j_*(*v*) (*e.g.*, arrows of Fig. 2). A parsimony is then some choice of exactly one box in this graph for each internal node of the phylogeny in such a way that they form an underlying directed spanning subtree in this graph, in terms of the arrows which join the boxes. This spanning subtree will have the same topology as the phylogeny, in fact, and the choice of box will correspond to the choice of ancestral state of the corresponding internal nodes of this phylogeny. In Fig. 2, for example, the spanning subtree *s*_low_(*X*)→*s*_low_(*Y*)→*s*_low_(*Z*) corresponds to the parsimony where weight is low in all ancestors, and *s*_low_(*X*)→*s*_high_(*Y*)→*s*_high_(*Z*) corresponds to the parsimony where high weight was ancestral in *Y*, and decreased in the Jaguarundi. Implicitly the leaves are also included in these spanning subtrees, but since there is no choice of state for extant species, they are left out for simplicity — see [8] for another example of this graph structure, with the leaves included. Notice, from Fig. 2 that there is a third solution *s*_high_(*X*)→*s*_high_(*Y*)→*s*_high_(*Z*) corresponding to a parsimony where high weight was ancestral (in *X*) to all species here, and then decreased in both the Pallas cat and the Jaguarundi. Note that for internal nodes *u* other than the root, that a state *i* ∈ Σ for which *s_i_*(*u*) is *not* minimized can appear in a solution, *e.g., s*_low_(*Y*). In general, for |Σ| = *k*, the procedure for building this graph structure (*e.g.*, in Fig. 2) would take time *O*(*nk*^2^), since at each node *u*, each of the *k* values *s_i_*(*u*) is derived from *O*(*k*) values at the (amortized constant number of) children of *u*.

Note that there can be *O*(*k*^*n*–1^) parsimonies in general. For example, consider the extreme case where all costs in *δ* are 0. For each internal node *v* and its parent u in the phylogeny, the set *s_i_*(*v*) of *k* boxes form a complete bipartite subgraph with the set *s_j_*(*u*) of *k* boxes in the corresponding graph structure. Any choice of *k* states in each of the *n* – 1 internal nodes (along with the unique choice for each leaf) would form a spanning subtree, giving rise to *k*^*n*–1^ parsimonies. In the general (less extreme) case, we first reduce the set of states for each internal node to those in a connected component of the graph containing a minimum score root state, as only those have the potential to be in a spanning subtree. For example, in Fig. 2, such (sets of) states are *s*_low_(*u*) and *s*_high_(*u*) for *u* ∈ {*X, Y, Z*}. Viewing the combinations of each possible choice of state for each internal node as elements of a Cartesian product (a subset of {1,…, *k*}^*n*–1^), a standard (*e.g.*, based on the left-hand rule) depth-first traversal (DFT) of this graph structure would visit the elements in *lexicographical order.* However, it is possible to visit the Cartesian product of a set in *Gray code* order [32], so that each consecutive element in the visit differs by exactly one co-ordinate. Visiting the graph structure in this manner allows for constant-time updates between consecutive potential combinations. That is, if a combination forms a spanning subtree, then the next combination, differing only by co-ordinate *u* (a node in the tree) updates only the edge to the parent and to the (amortized constant number of) children of *u* in this spanning subtree. It then suffices to check if this update is also a spanning subtree in order to verify if the current combination also forms a spanning subtree. While this graph structure (Fig. 2) is mentioned [8], how to efficiently enumerate all its parsimonies is rarely discussed (that we know of). On the other hand, since having all parsimonies is central to our approach, the above Gray code enumeration strategy is implemented into `parcours`.

## 3 Correlation

We want to determine correlated evolution among pairs of characters. Suppose we have a second character, dental profile (number of teeth), which can have states from {28, 30, unknown}. Here, the dental profile is “fewer” (28) only in the Pallas cat as depicted in Fig. 3a, and we want to understand how this is correlated to the weight of Fig. 1b.

**Figure 3:**
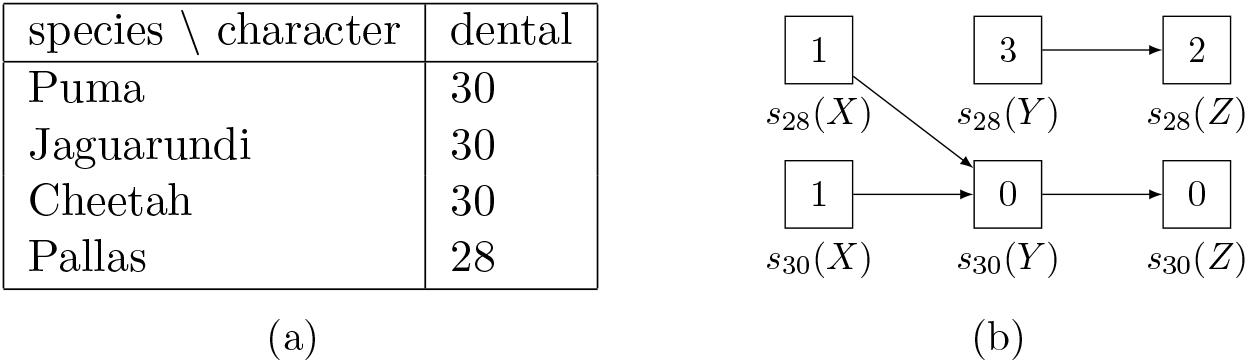
The (a) extant state of character dental in four species and (b) the structure resulting from Sankoff’s algorithm on these extant states along with the phylogeny of Fig. 1a and cost of Fig. 1c. Values *s*_2_ have been omitted for compactness, since they are all 8, like in Fig. 2.

The idea is that we first construct likely hypotheses for what ancestral states each character *α* might have, namely the set *P*(*α*) of all small parsimonies. Then we determine if there are any correlated changes in state of pairs of characters along branches of the phylogenetic tree given their sets of parsimonies. While a drawback of parsimony is potentially many solutions, without any a priori knowledge of ancestral state, choosing one solution (or a subset of solutions) is an arbitrary decision. Since such a decision could potentially bias any downstream analysis, our approach attempts to avoid this, in using the entire set of parsimonies. Let *i*→*j* be a change in state (for some pair *i, j* of states) of character *α*. Let *p*|_*i*→*j*_ be the *multiset* (even though each element will have multiplicity 0 or 1) of *branches* in the phylogenetic tree where change *i*→*j* occurs in parsimony *p* ∈ *P*(*α*). The *consensus map C_α_*(*i*→*j*) for change *i*→*j* of character *α* is then the multiset resulting from

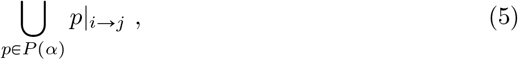

in preserving the information of *all* parsimonies. The idea is that the consensus map captures the area of the tree (a set of branches), where this type of change occurs. Note that being a multiset is important, as the same change from different parsimonies may occur on the same branch of the tree, and we want to count this. The *correlation* between some change *i*→*j* of character *α* and some change *i*′→*j*′ of a character *β* is then the *weighted Jaccard index* [24], also known as the Ružička index ([39])

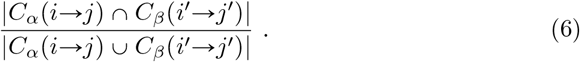

The weighted Jaccard index measures the degree to which the respective consensus maps overlap, taking into account multiplicities on the branches.

For example, given the characters weight (Fig. 1b) and dental profile (Fig. 3a), we compute the correlation of the change low→high (in weight) and 28→30 (in dental profile) in the phylogeny of Fig. 1a (according to cost of Fig. 1c) as follows. Character weight has the set *P*(weight) = {*p*_1_,*p*_2_,*p*_3_} of three parsimonies, with *p*_1_|_low→high_ = {*Z*→Puma,*Y*→Cheetah}, *p*_2_|_low→high_ = {*X*→*Y*}, and *p*_3_|_low→high_ = ⊘. It follows from Eq. 5 that the consensus map *C*_weight_ (low→high) = {*Z*→Puma, *Y*→Cheetah, *X*→*Y*}. By inspecting Fig. 3b, character dental has the set *P*(dental) = {*q*_1_, *q*_2_} of two parsimonies, with *q*_1_|_28→30_ = {*X*→*Y*} and *q*_2_|28→30 = ⊘, hence *C*_dental_(28→30) = {*X*→*Y*}. It then follows from Eq. 6 that the correlation of this co-event is 1/3 « 0.33. The correlation of the co-event high→low and 30→28 in weight and dental is also 1/3, while the correlation of low→high in weight and 30→28 in dental (and high→low in weight and 28→30 in dental) is 0, because there is no overlap between sets *p*|_low→high_ and *q*|_30→28_ for any pair of parsimonies *p* ∈ *P*(weight) and *q* ∈ *P*(dental) (and vice versa). For completeness, the correlation of any combination of event involving an unknown state in one character with any other event in the other character is 0 because none of these events happen in any parsimony of weight or dental. We use the weighted Jaccard index because it measures the amount of event co-occurrence, normalized by the amount of independent occurrences of either event in the set of all parsimonies, taking multiplicities into account. If one wanted focus on just the events on the different branches (without multiplicity), one could use the unweighted Jaccard index, which is Eq. 6 where all multisets are treated as *sets* (all non-zero multiplicities cast to 1).

In a similar approach [33], albeit in a maximum likelihood setting where there are branch lengths, the authors map the probabilities of gain and loss to each branch of the phylogeny in the most likely ancestral assignment. Correlation between such events in a pair of characters is then assessed by computing the mutual information between the associated sets of probabilities. Our approach could be viewed as a discrete analog of this.

## 4 Significant Correlation

The correlation between two changes (in two characters), presented in Section 3, is the degree of overlap between their consensus maps. However, we want to know if this correlation is *significant*, that is, if it is higher than expected by chance. A consensus map *C_α_*(*i→j*) for change *i→j* in character *α* is constructed from the set *P*(*α*) of small parsimonies, each of which represents a (smallest) number of particular changes on the tree. Here, the null hypothesis posits that these particular changes happened randomly on the tree. A “random” consensus map for change *i→j* of character α is then obtained assuming that the changes, in each *p* ∈ *P*(*α*) of the parsimonies it is constructed from, happened randomly on the tree. To determine if a correlation between some change *i*→*j* in character *α* and some change *i*′→*j*′ in character *β* is significant, we need to determine if the probability of obtaining this correlation is significantly higher than the correlation of a pair of random consensus maps for *i*→*j* and *i*′→*j*′. To do this, we need to compute, for change *i*→*j* (and *i*′→*j*′, respectively), the collection 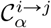 (and 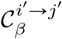, respectively) of consensus maps obtained from all possible combinations of ways in each *p* ∈ *P*(*α*), of distributing the changes in *p* on the tree. A (null) distribution of correlation is then obtained by computing the correlation for each possible pair of consensus maps from collections 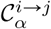 and 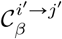. A correlation between *i*→*j* and *i’→j’* is then deemed significant if the probability of obtaining it is sufficiently extreme in this distribution.

We continue the example of the phylogenetic tree of Fig. 1a and characters weight (Fig. 1b) and dental profile (Fig. 3a) to illustrate the notion of significant correlation. For compactness, we relabel Puma, Jaguarundi, Cheetah and Pallas Cat, species *A, B, C* and *D*, respectively, as depicted in Fig. 4a. We also relabel weight and dental profile, characters *α* and *β*, respectively (Fig. 4b). For further compactness, we refer to a branch according to its *descendant* species, and so, *e.g.*, the branch from *Y* to *C* would be referred to as *C*. For even further compactness, since our characters *α* and *β* are binary, for a given branch, *e.g., Y*, we use the upper case *Y* to refer to a change of state low→high or 28→30 (an increase) for their respective characters *α* and *β* on this branch, and the lower case *y* to refer to change high→low or 30→28 (a decrease) on this branch, respectively, when the context is clear.

**Figure 4:**
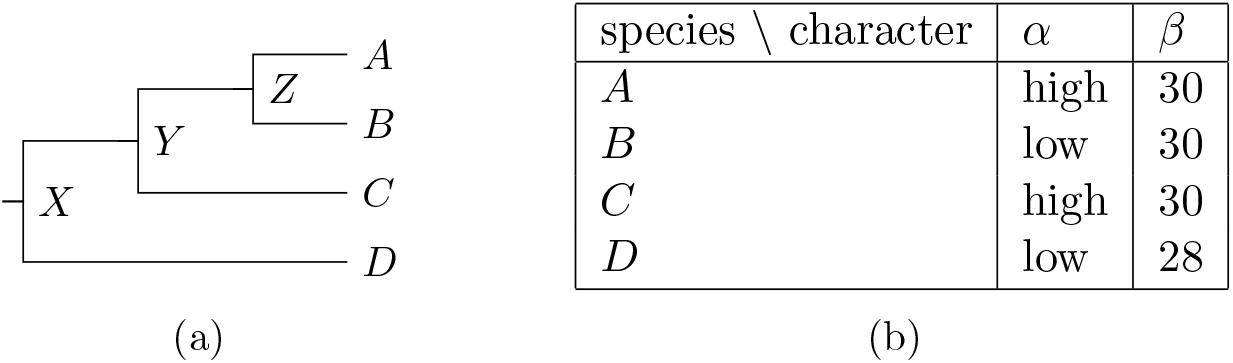
A (a) phylogeny and (b) the extant states of characters *α* and *β* in four species *A, B, C* and *D*.

For character *α* (weight), the set *P*(*α*) is then *p*_1_ = {*A, C*}, *p*_2_ = {*b, d*} and *p*_3_ = {*Y, b*}, and for character *β* (dental profile), the set *P*(*β*) is then *q*_1_ = {*Y*} and *q*_2_ = {*d*}. It follows that the consensus maps *C_α_*(low→high) and *C_α_*(high→low), for the changes low→high and high→low in character *α*, respectively, are

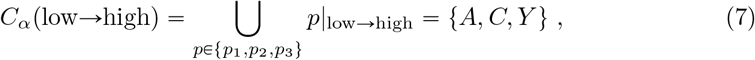

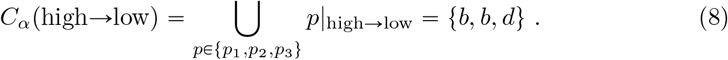

The consensus maps for the changes 28→30 and 30→28 in character *β* are then *C_β_*(28→30) = *q*_1_ = {*Y*} and *C_β_*(30→28) = *q*_2_ = {*d*}, respectively. The correlation of a change low→high in character *α* with 28→30 in character *β* is then

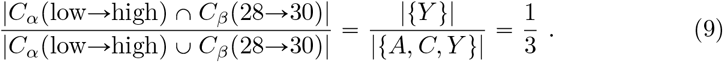

To determine if this correlation is significant, we need to compute the null distribution — the correlation between each pair from (the Cartesian product of) collections 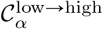 and 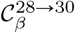. Consensus map *C_α_*(low→high) comes from *P*(*α*) = {*p*_1_, *p*_2_, *p*_3_}, so we need to compute all possible combinations of ways that the events of each of *p*_1_, *p*_2_ and *p*_3_ can happen on the tree, and then take a consensus map from each member of the (Cartesian) product of these sets of possible ways to obtain 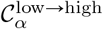. For some alphabet Σ, let *t*(Σ) = {*i*→*j*| *i, j* ∈ Σ, *j* ≠ *i*}, that is, the set of possible changes (or transitions) between any pair of characters from Σ. For example, the alphabet of character *α* is Σ_*α*_ = {low, high}, so *t*(Σ_*α*_) = {low→high, high→low}. Let *e^α^*(*p*) then be the number of instances of each type of change (or event) from *t*(Σ_*α*_) in parsimony *p* ∈ *P*(*α*) for character *α* — effectively a multiset — expressed as a tuple (in some fixed ordering, *e.g.*, lexicographic, on Σ_*α*_). For example, where Σ_*α*_ = {low, high}, *e^α^*(*p*_2_) = (2,0), for 2 low→high events and 0 high→low events, while *e^α^*(*p*_2_) = (0, 2) and *e^α^*(*p*_3_) = (1,1). Now, let 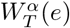 be the set of ways that some multiset *e* of events (from set *t*(Σ_*α*_) of possible events for character *α*) can happen on a tree *T*. For example, in phylogeny (call it *T*) of Fig. 4a, 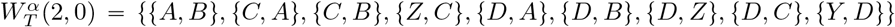 — the set of ways that 2 low→high events (and 0 high→low events) in character *α* could happen on *T*. Note that *p*_1_, *p*_2_ and *p*_3_ are each just one of the possible ways from 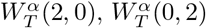 and 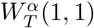, respectively. The collection 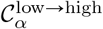 is then

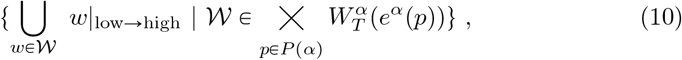

which is effectively the consensus map *C_α_*(low→high) (Eq.5) for change low→high of character *α* from each element (one of the of possible combinations of ways that the events of each *p* ∈ *P*(*α*) can happen on the tree) of the Cartesian product 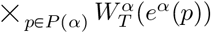. The remaining detail lies in how we compute 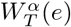 for some multiset e of events for character *α* on a tree *T*.

Computing 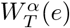 for some multiset *e* of events in character *α* on a tree *T* tree is done using a dynamic programming approach which combines the sets of possible ways the events can happen on subtrees of *T* in a bottom-up fashion. This is similar to the computation of *W_F_*(*n,m*|*b*) from [29], hence the same *W* notation.^2^ More precisely, for some node *u* in tree *T*, 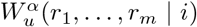 is the set of ways of having *r_t_* events of type *t* (*e.g.*, a low→high event), for *t* ∈ {1,…, *m*}, given that the state (from alphabet Σ_*α*_) of node *u* is i. That is, the *r*_1_,…,*r_m_* are just the elements of the tuple representing some multiset *e* of events from *t*(Σ_*α*_) in character *α, i.e., m* = |*t*(Σ_*α*_)|. The 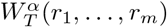 from above is then

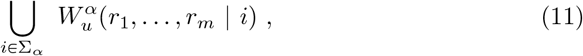

where *u* is the root of tree *T*. Since this is computed using a bottom-up dynamic programming procedure, the base case for each leaf node ℓ of tree *T* is

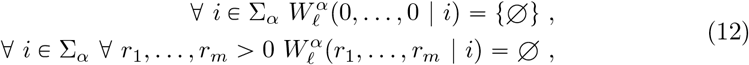

where ⊘ is the empty set. The recursive step for internal node *u* with the set *V* = {*v*_1_,…, *v_n_*} children is then

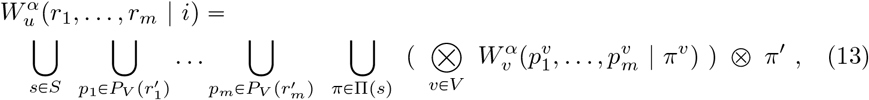

where

- *S* is the number of multisets of size *n* taken from Σ_*α*_, *i.e.*, the number of ways to distribute events on the branches below *u* (note that *i→i* is the non-event);
- 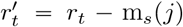, if *t* represents transition (event) *i*→*j* and *m_s_*(*j*) is the *multiplicity* of element *j* in multiset *s* (o.w., 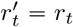);
- 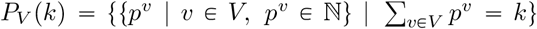 is the number of ways to partition *k* objects into the (*n*) bins *V*, i.e., the number of ways to distribute the remaining events to the subtrees below *u*;
- Π(*s*) is the set of *unique* permutations of multiset *s*;
- 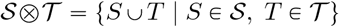, which is somewhat like a Cartesian product, but it returns a set of sets, rather than a set of tuples;
- *π^v^* is the *v*-th element of permutation *π*; and
- *π*^1^ = {{*v*: *i*→*j* | *v* ∈ *V, j* = *π^v^* ≠ *i*}} is the set of changes *v*: *i*→*j* from some state *i* to some other state *j* ≠ *i* happening on the branch leading to node v from its parent (node *u*).

Note that this recursive step at node *u* is just computing all possible ways to have any *v*: *i*→*j* event on any combination of each branch leading to some child *v* ∈ *V*, for all combinations of possible ways 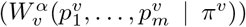 events can happen in the subtree rooted at each node *v* ∈ *V*. At any given step, the set, *e.g.*, 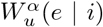 of ways some multiset *e* of events in character *α* can happen on tree *T*, given that the state of *α* at node *u* is *i*, is then just a set of sets of these *v*: *i*→*j* events. For example, if character *α* represents weight from above, then the set 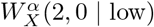 of ways that 2 low→high events and 0 high→low events can happen on the tree *T* of Fig. 4a, given that the state of character *α* at node *X* in *T* is low, is

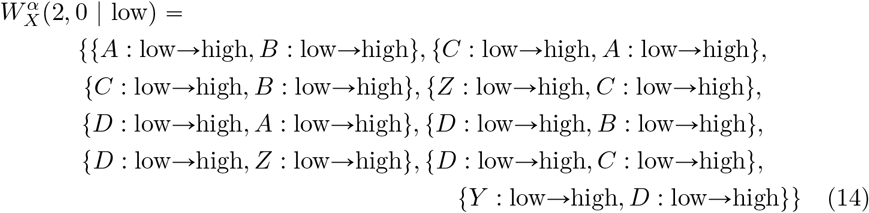

or {{*A, B*}, {*C, A*}, {*C, B*}, {*Z, C*}, {*D, A*}, {*D, B*}, {*D, Z*}, {*D, C*}, {*Y, D*}} in our shorthand devised for binary characters (see Eq. 55 of Appendix A).

We now detail the computation of the correlation between each pair from the Cartesian product of collections 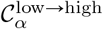 and 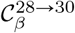 for the example depicted in Fig. 4. To compute 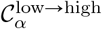, based on Eq. 10, we first compute 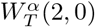, 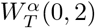 and 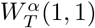 according to Eq. 11 — the ways that the events of each of the respective parsimonies *p*_i_, *p*_2_ and *p*_3_ for character *α* can happen on *T* of Fig. 4a, as follows

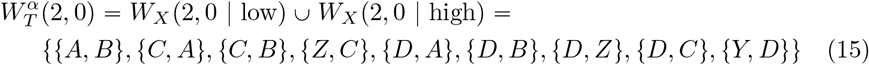

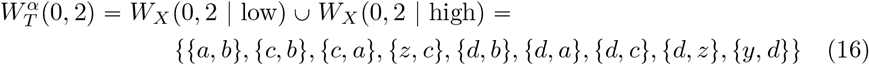

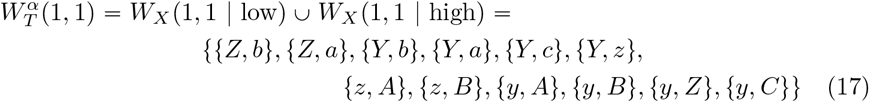

Note that the details of computing 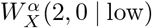 (Eq. 55), 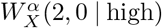 (Eq. 58), 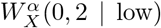 (Eq. 53), 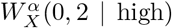 (Eq. 56), 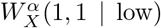 (Eq. 54) and 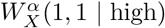 (Eq. 57) are found in Appendix A. Collection 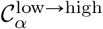 is then the (9 · 9 · 12 = 972) sets resulting from the Cartesian product of 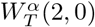, 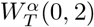 and 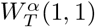 (from Eqs. 15, 16 and 17, respectively) computed according to Eq. 10. The first two elements and the last element of 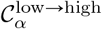 are as follows

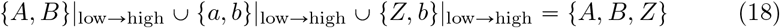

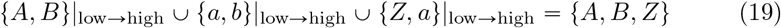

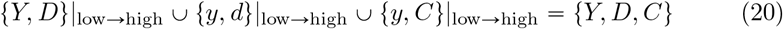

To compute 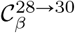, we first compute 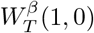 and 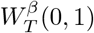 according to Eq. 11 — the ways that the events of each of the respective parsimonies *q*_1_ and *q*_2_ for character *β* can happen on *T* of Fig. 4a as follows

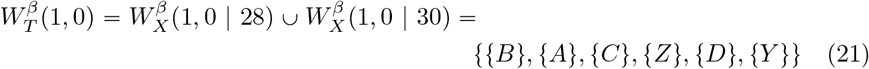

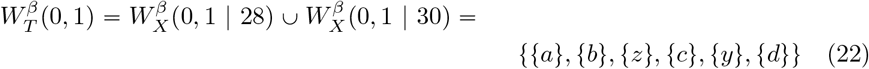

Note that the details of computing 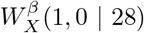 (Eq. 72), 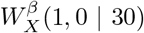 (Eq. 74), 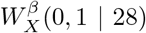 (Eq. 71) and 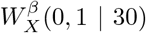 (Eq. 73) are found in Appendix A. Collection 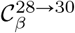 is then the (6 · 6 = 36) sets resulting from the Cartesian product of 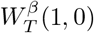 and 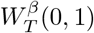 (from Eqs 21 and 22, respectively) computed according to Eq. 10. The first two elements and the last element of 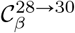 are as follows

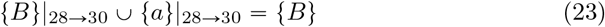

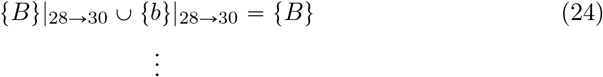

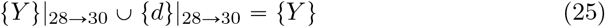

Finally, the set of correlations (of size 972 · 36 = 34992) obtained by computing the correlation (according to Eq. 6) of each pair from the Cartesian product of the collections 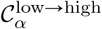 and 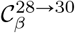 is computed. The first two correlations and last correlation of this set are as follows

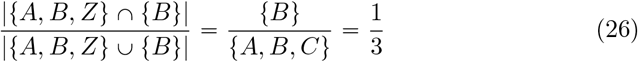

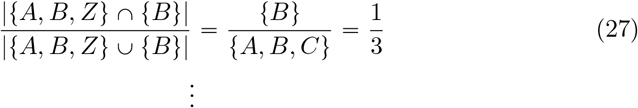

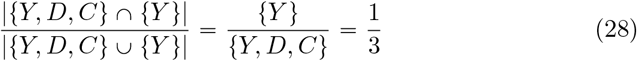

Of these 34992 correlations, 16092 of them are 1/3, and 18900 of them are 0. This distribution has a mean of *μ* = 0.153 and a standard deviation of *σ* = 0.166. It follows that the probability of obtaining a correlation as extreme as 1 /3 in this (null) distribution of correlation is 0.46 (*p*-value), and 1/3 is 1.084 *σ* from the mean (*z*-score), hence this correlation is far from being significant by standard conventions (*p* < 0.05 or *z* ≽ 2).

## 5 Experimental Analysis

To test our approach, we used a set of vocalization and morphological data from the subfamily Pantherinae, commonly known as the big cats. We chose this clade in our analyses because their vocalization data is best documented among Felidae. Pantherinae consists of six species and two genera: *Panthera* (lions, leopards, jaguars, tigers and snow leopards) and *Neofelis* (clouded leopards) (Fig. 5). The set of extant states are for 10 of the vocalizations documented in [46], and 9 morphological characteristics compiled from the various sources within [5], see Table 1. The vocalizations had states {absent, present, unknown}, while the morphological characteristics where binned into ranks starting from 1 (largest, or longest, etc.), to 2, to 3, etc. Finally, the cost for changing to a different state is 1, and to an *unknown* state is ∞ (*e.g.*, Fig. 1c). Since unknown states are artifacts of the collection process, by assigning a high evolutionary cost to the changes from any unknown to some known state (which makes little evolutionary sense), we mitigate the propagation of any unknown state to ancestral nodes in any parsimony, whenever some known state (*e.g.*, low or high) can explain the data. For example, in the instance of Fig. 1, if the weight was instead unknown in the Jaguarundi and Pallas cat, then it would only have the unique parsimony where all ancestors *X, Y* and *Z* have the high state. This is why we use the approach of [40] (instead of, *e.g*., [20]), because we need these transition-specific costs.

**Figure 5:**
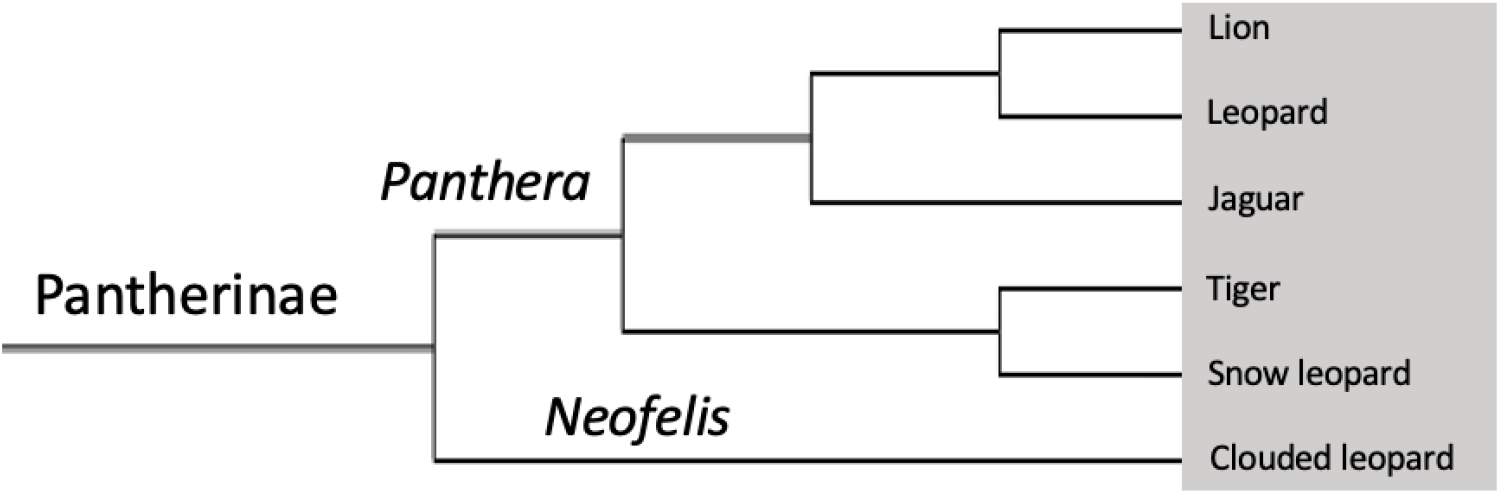
The subfamily Pantherinae consists of six species and two genera: *Panthera* (lions, leopards, jaguars, tigers and snow leopards) and *Neofelis* (clouded leopards). Phylogeny adapted from [26] and [28, 19].

**Table 1:** Vocalizations and morphological characteristics used in the experiment.

| Vocalizations | Morphological characteristics |
| --- | --- |
| spit | gestation time |
| hiss | skull width (SW) |
| growl | skull length (SL) |
| snarl | body length (BL) |
| prusten | dental profile |
| puff | weight |
| mew | skull length / skull width ratio (SL/SW) |
| main call | body length / skull length ratio (BL/SL) |
| roaring sequence | chromosome number |
| grunt |  |

Our approach and resulting implementation into the software tool `parcours` begins with the efficient computation of all parsimonies described in Sec. 2, followed by the computation of correlation from these parsimonies, as described in Sec. 3. Finally, for each of these correlations, the probability of obtaining a correlation as extreme (or more) by chance is computed according to Sec. 4. Since `parcours` starts with the same input as Sankoff’s algorithm (*e.g.*, Fig. 1), we input the phylogenetic tree and extant states of the 19 characters and the Pantherinae phylogeny mentioned above. The cost matrix is computed automatically by `parcours`, this being the default, when no specific cost matrix is provided. From this, `parcours` returns the correlations of all possible events between all pairs of characters according to Eq. 6, and then computes the probability obtaining a correlation this extreme by chance according to Sec. 4.

## 6 Results and Discussion

We used two measures of significance. In a normal distribution, a standard *p* value of 0.05 is equivalent to a *z* score of two or more. However, because the distributions here are computed from a (large, yet finite) number of ways some sets of events may occur on a tree, they are discrete, not normal distributions. As such, we defined significance using either *p* < 0.05 and/or *z* score of two or more. Table 2 reports all correlations found by `parcours` where either *p* < 0.05 or *z* ≽ 2. We examined all such significant co-occurrences to assess if they were consistent with known biological data as a way to evaluate `parcours`.

**Table 2:**
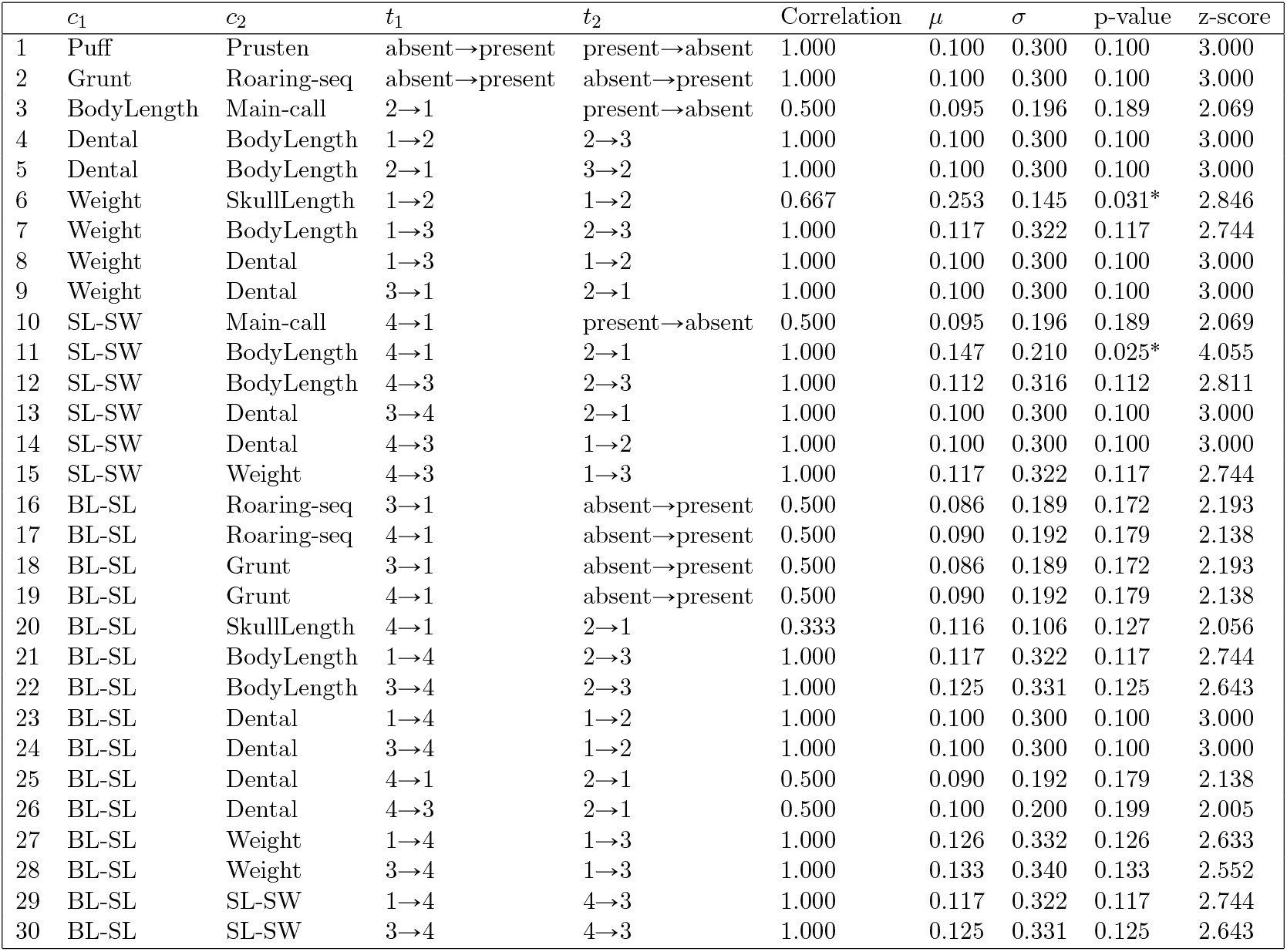
Of the 61 pairs of events with nonzero correlation, the 30 which are significantly correlated, as judged by a p-value < 0.05 or a *z*-score ≽ 2. Each row is correlation between a pair *t*_1_ and *t*_2_ of events in characters *c*_1_ and *c*_2_, respectively. The *μ* and *σ* are the mean and standard deviation of the corresponding null distribution, respectively. The p-value is the probability of obtaining a correlation this extreme in the null distribution, while the *z*-score is the number of standard deviations (*σ*) this probability (*p*) is from the mean (*μ*).

| | $c_1$ | $c_2$ | $t_1$ | $t_2$ | Correlation | $\mu$ | $\sigma$ | p-value | z-score |
| --- | --- | --- | --- | --- | --- | --- | --- | --- | --- |
| 1 | Puff | Prusten | absent→present | present→absent | 1.000 | 0.100 | 0.300 | 0.100 | 3.000 |
| 2 | Grunt | Roaring-seq | absent→present | absent→present | 1.000 | 0.100 | 0.300 | 0.100 | 3.000 |
| 3 | BodyLength | Main-call | 2→1 | present→absent | 0.500 | 0.095 | 0.196 | 0.189 | 2.069 |
| 4 | Dental | BodyLength | 1→2 | 2→3 | 1.000 | 0.100 | 0.300 | 0.100 | 3.000 |
| 5 | Dental | BodyLength | 2→1 | 3→2 | 1.000 | 0.100 | 0.300 | 0.100 | 3.000 |
| 6 | Weight | SkullLength | 1→2 | 1→2 | 0.667 | 0.253 | 0.145 | 0.031* | 2.846 |
| 7 | Weight | BodyLength | 1→3 | 2→3 | 1.000 | 0.117 | 0.322 | 0.117 | 2.744 |
| 8 | Weight | Dental | 1→3 | 1→2 | 1.000 | 0.100 | 0.300 | 0.100 | 3.000 |
| 9 | Weight | Dental | 3→1 | 2→1 | 1.000 | 0.100 | 0.300 | 0.100 | 3.000 |
| 10 | SL-SW | Main-call | 4→1 | present→absent | 0.500 | 0.095 | 0.196 | 0.189 | 2.069 |
| 11 | SL-SW | BodyLength | 4→1 | 2→1 | 1.000 | 0.147 | 0.210 | 0.025* | 4.055 |
| 12 | SL-SW | BodyLength | 4→3 | 2→3 | 1.000 | 0.112 | 0.316 | 0.112 | 2.811 |
| 13 | SL-SW | Dental | 3→4 | 2→1 | 1.000 | 0.100 | 0.300 | 0.100 | 3.000 |
| 14 | SL-SW | Dental | 4→3 | 1→2 | 1.000 | 0.100 | 0.300 | 0.100 | 3.000 |
| 15 | SL-SW | Weight | 4→3 | 1→3 | 1.000 | 0.117 | 0.322 | 0.117 | 2.744 |
| 16 | BL-SL | Roaring-seq | 3→1 | absent→present | 0.500 | 0.086 | 0.189 | 0.172 | 2.193 |
| 17 | BL-SL | Roaring-seq | 4→1 | absent→present | 0.500 | 0.090 | 0.192 | 0.179 | 2.138 |
| 18 | BL-SL | Grunt | 3→1 | absent→present | 0.500 | 0.086 | 0.189 | 0.172 | 2.193 |
| 19 | BL-SL | Grunt | 4→1 | absent→present | 0.500 | 0.090 | 0.192 | 0.179 | 2.138 |
| 20 | BL-SL | SkullLength | 4→1 | 2→1 | 0.333 | 0.116 | 0.106 | 0.127 | 2.056 |
| 21 | BL-SL | BodyLength | 1→4 | 2→3 | 1.000 | 0.117 | 0.322 | 0.117 | 2.744 |
| 22 | BL-SL | BodyLength | 3→4 | 2→3 | 1.000 | 0.125 | 0.331 | 0.125 | 2.643 |
| 23 | BL-SL | Dental | 1→4 | 1→2 | 1.000 | 0.100 | 0.300 | 0.100 | 3.000 |
| 24 | BL-SL | Dental | 3→4 | 1→2 | 1.000 | 0.100 | 0.300 | 0.100 | 3.000 |
| 25 | BL-SL | Dental | 4→1 | 2→1 | 0.500 | 0.090 | 0.192 | 0.179 | 2.138 |
| 26 | BL-SL | Dental | 4→3 | 2→1 | 0.500 | 0.100 | 0.200 | 0.199 | 2.005 |
| 27 | BL-SL | Weight | 1→4 | 1→3 | 1.000 | 0.126 | 0.332 | 0.126 | 2.633 |
| 28 | BL-SL | Weight | 3→4 | 1→3 | 1.000 | 0.133 | 0.340 | 0.133 | 2.552 |
| 29 | BL-SL | SL-SW | 1→4 | 4→3 | 1.000 | 0.117 | 0.322 | 0.117 | 2.744 |
| 30 | BL-SL | SL-SW | 3→4 | 4→3 | 1.000 | 0.125 | 0.331 | 0.125 | 2.643 |

Out of the 30 significant correlations detected by `parcours`, 12 were between traits associated with overall body size such as weight and body length (BL). This is not surprising given that many features like body weight and body length have allometric relationships [3]. Significant co-occurrences within Pantherinae include a decrease in both weight and skull length (SL) (*p* = 0.03) and an increase in both the skull length / skull width (SL/SW) ratio and body length (*p* = 0.025). For example, these significant co-occurrence were detected in the lineage leading to the African leopard and African lion, respectively. These findings are consistent with the observation that (a) African leopards are the smaller in both weight and skull length than either of its sister taxa, the lions and jaguars and (b) lions have the largest body length and SL/SW ratio of any member of Pantherinae.

Prusten and puff are two types of friendly close-range vocalizations. Felid species have only one type of friendly call in their repertoire as the prusten and puff are functionally equivalent [46]. Prusten is found in all members of Pantherinae except for lions and leopards which instead have a puff [37]. A significant co-occurrence of a loss of prusten and a gain of puff (*z* = 3.0) along the lineage leading to lions and leopards, is consistent with the observation that prusten is most likely the ancestral friendly call type in Pantherinae which was replaced by a puff in lions and leopards.

The dental formula (DF) is defined as the total number of each type of tooth in both the upper and lower jaw. Within Pantherinae, all species have a DF of 30 with the exception of the Indochinese Clouded Leopard (*Neofelis nebulosa*) which has a DF of 28-30. Clouded leopards are the smallest member of Pantherinae in terms of weight and overall body length. Weighing an average of 17 kg and 88 cm in length, the clouded leopards are mixed prey feeders, adapted to taking both large and small prey. All other members of Pantherinae are large prey feeders with relatively wide muzzles [31]. This is consistent with the observation that a decrease in SL/SW ratio co-occurs with an increase in dental formula and vice versa (*z* = 3.0). The direct co-occurrence of body length with dental formula and weight with dental formula (*z* = 3.0) is also consistent with the fact that the clouded leopard is the smallest felid in Pantherinae and the only felid with a dental formula less than 30.

A roaring sequence is characterized by one to two soft moans followed by full throated roars and a series of grunts of diminishing amplitude [44]. There was a significant co-occurrence between an increase in BL/SL ratio from a medium sized to a large bodied *Panthera* species and the gain of this call along the lineage leading to lions, leopards and jaguars (*z* = 2.2). This is consistent with the fact that the roaring sequence is known only those felid species. While grunts are most often used as part of a roaring sequence, grunts can also be used alone in certain situations such as females calling for cubs [46]. Not surprisingly, the gain of a grunt also significantly co-occurs with an increase in BL/SL ratio along the same lineage (*z* = 2.2). Finally, there is significant association between the gain of a grunt and a gain in the roaring sequence (*z* = 3.0), also along this same lineage. This is consistent with the observation that the grunt is an integral part of the roaring sequence supports their non-random co-occurrence.

The main call is defined as a high intensity form of a mew. Mews are the predominant vocalization of felids but have low to medium intensities. Main calls are found in all felids except for lions [46]. We detected a significant co-occurrence in the increase in both SL/SW ratio and body length with the loss of a main call (*z* = 2.1) along the lineage leading to lions. This is consistent with main calls being absent in lions, and African lions being the largest felid within Pantherinae in terms of body length, and having the highest SL/SW ratio of all felids in this subfamily.

## 7 Conclusion

Understanding the evolution of complex traits such as vocalization and morphological characters remains a key challenge in evolutionary studies. In this work, we propose a new approach to detecting significantly correlated evolution among discrete characters in a phylogenetic tree by leveraging the information from *all* parsimonies — its implementation, `parcours`, efficiently automating this process in an easy-to-use and customizable way. We demonstrate its use on a set vocalizations and morphological characteristics in the Pantherinae subfamily.

While `parcours` already discovers trends in this data which are supported in the literature, possible refinements to the ranking of characteristics (such as dental profile), or the cost matrix (*e.g.*, making 4 → 1 more expensive than 2 → 1), could even further improve these results. While we used the weighted Jaccard index as correlation here, a full exploration of similarity indexes [48] is the subject of future work. Here, we consider the set of all parsimonies in this framework. However, another idea is to allow the user to place constraints on this set, based on a priori knowledge, such as certain ancestral states, or the least common ancestor of a given set of states. To assess significant correlation, we compute all possible ways some sets of events can happen on a tree. However, this can result in runtime issues, something which was encountered also in the method [29] which inspired the design of our approach. A potential solution could be to subsample such set of possible ways, allowing for a faster approximation of significant correlation. Finally, a future experimental direction is to reconstruct even more detailed information about felid vocalizations (*e.g.*, mean frequency, harmonics) from audio recordings. We have already devised a machine learning framework for automatically extracting many such acoustic features from sound recordings, and have some interesting preliminary results on the Pantherinae subfamily [1]. Using `parcours` to then automate the search for significant correlations among the many features extracted, along with morphological characters (*e.g.*, mean frequency and body size), will provide valuable insight into the evolution of their vocalizations in the context of morphology and perhaps even habitat.

## Acknowledgements

Research supported by Fairfield University Science Institute and Fredrickson Family Innovation Lab grants for SAB and MP; Georgia State University startup grant for MP. The authors would like to thank Steven Patterson for insightful discussions about the statistical aspects of this work.

## A Appendix

### Ways that two events in character *α* can happen in a tree

Given character *α* from Fig. 4b (Σ_*α*_ = {low, high}), we detail how to compute all possible ways that two events, from *t*(Σ_*α*_) = {low→high, high→low}, can happen on tree *T* of Fig. 4a, given either state low or high at the root (*X*) of the tree *T, i.e.*, 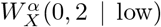 (Eq. 53), 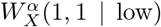 (Eq. 54), 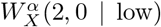 (Eq. 55), 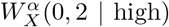 (Eq. 56), 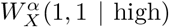 (Eq. 57) and 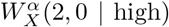 (Eq. 58). The base case(s) for the leaf nodes *A, B, C* and *D* of tree *T* is set according to Eqs. 12 as follows

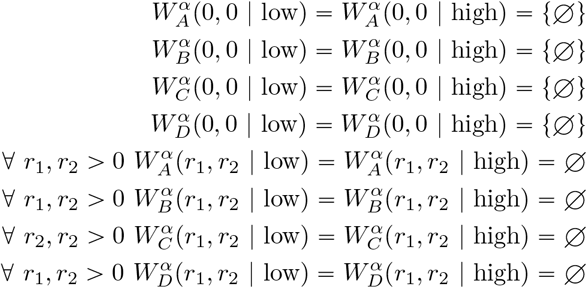

The recursive step(s) for each internal node *Z, Y* and finally *X* of tree *T* is then computed according to Eq. 13 as follows

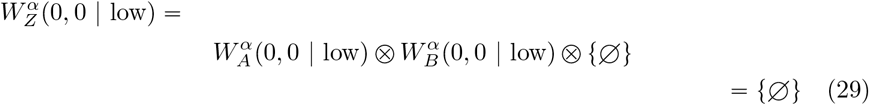

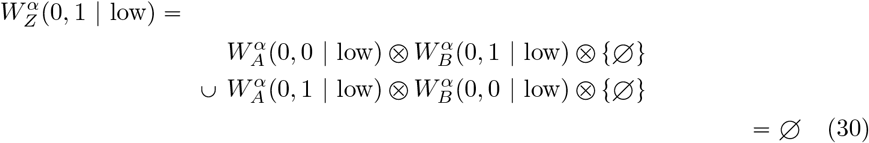

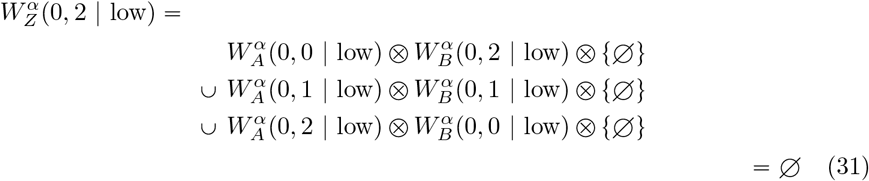

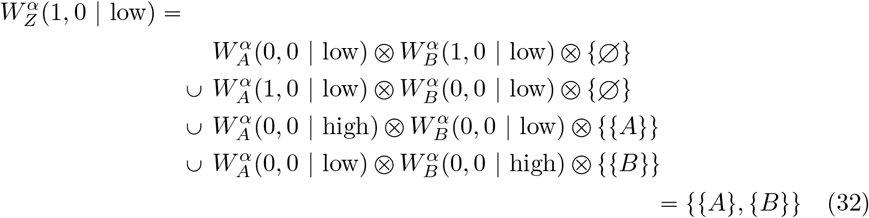

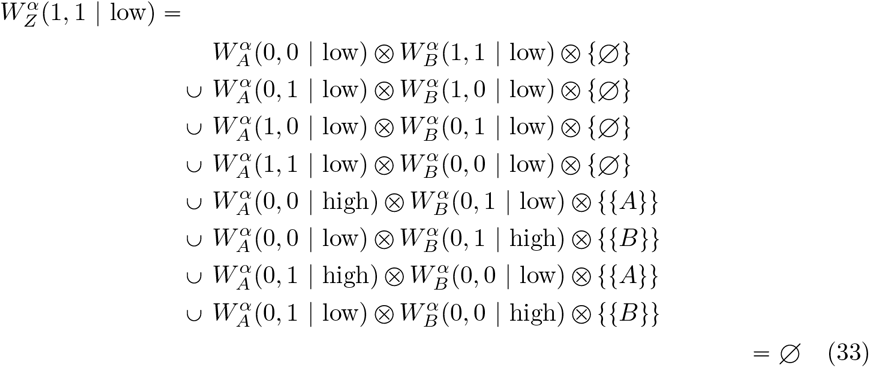

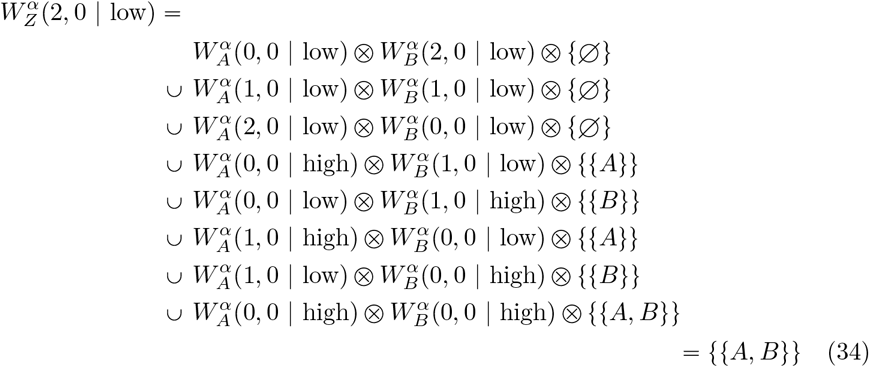

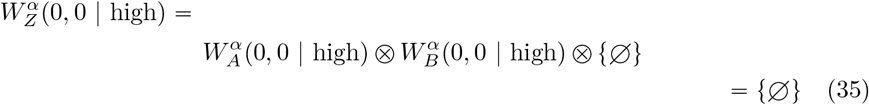

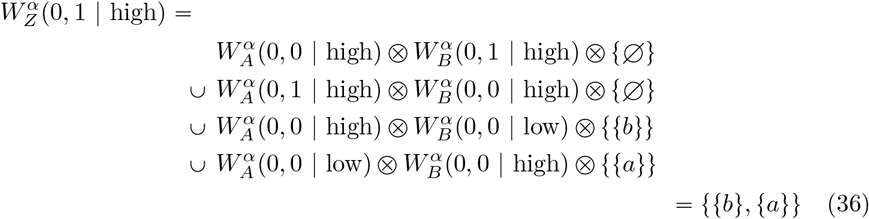

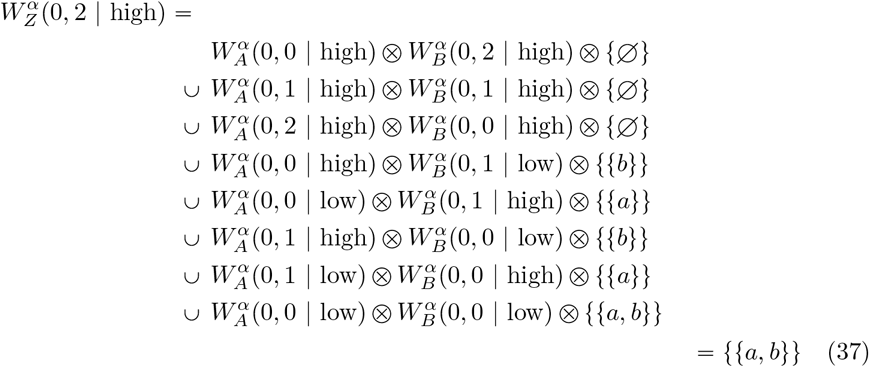

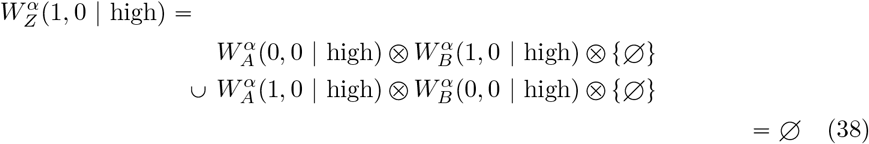

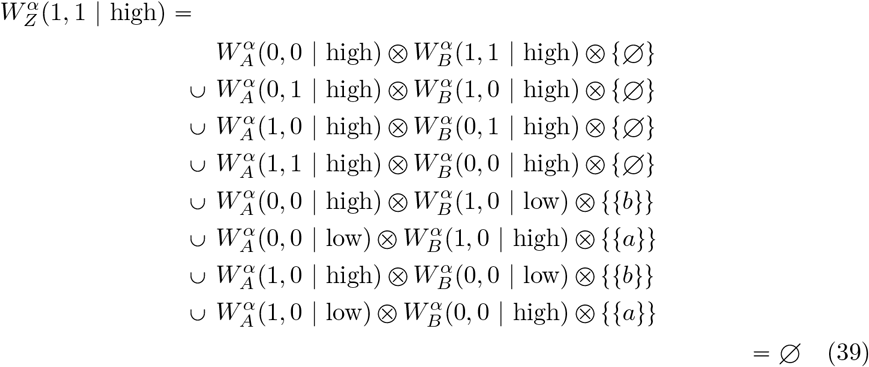

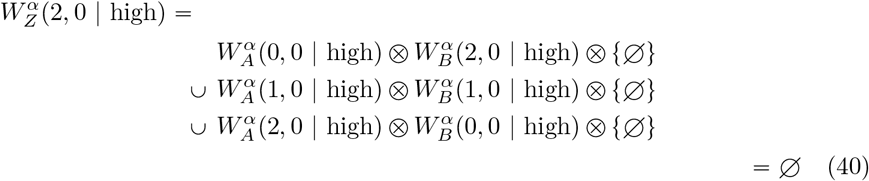

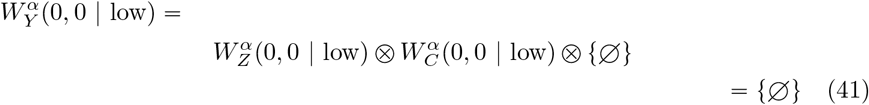

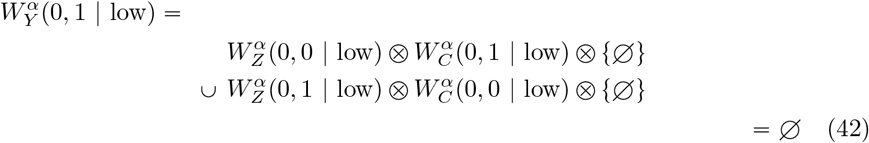

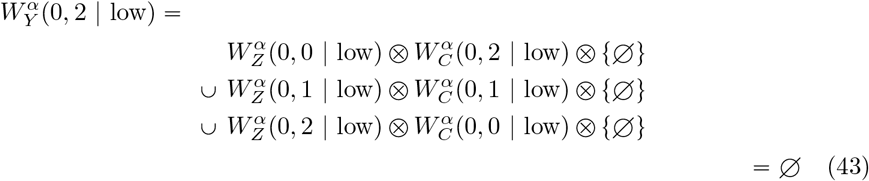

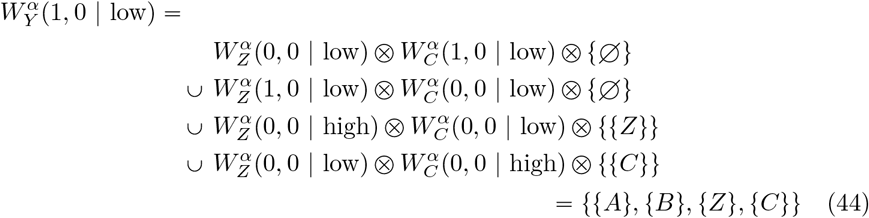

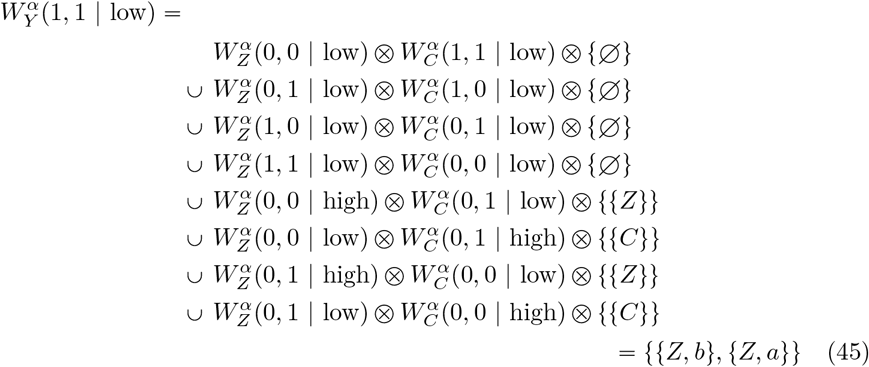

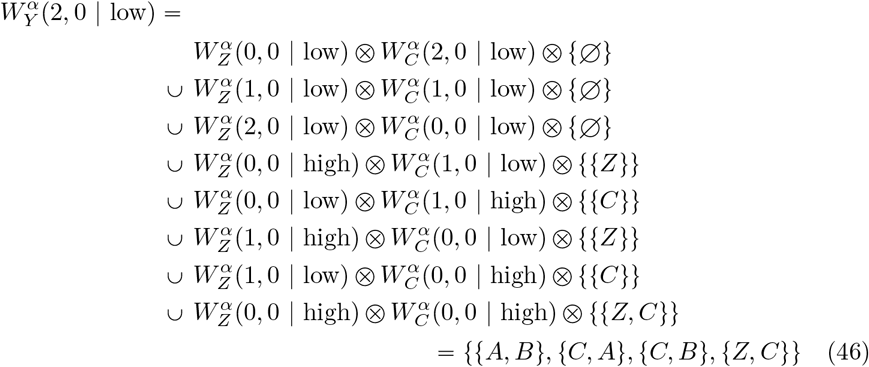

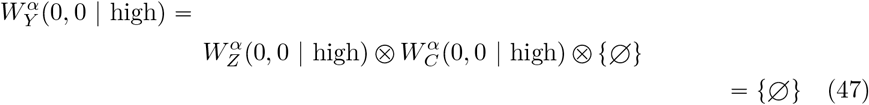

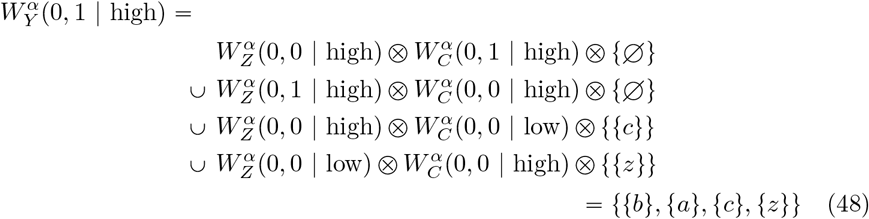

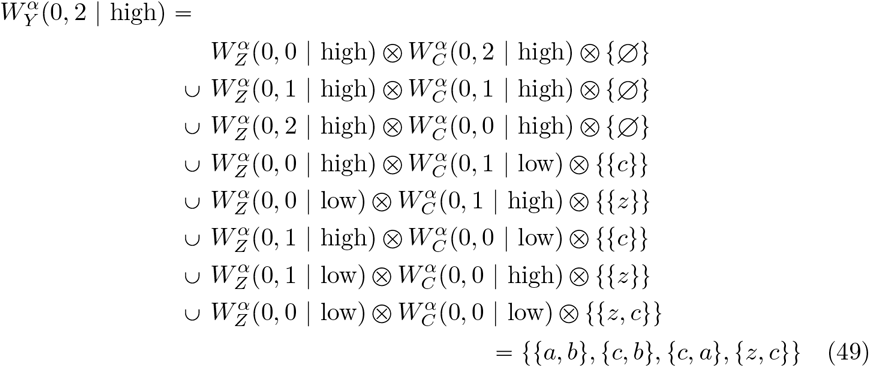

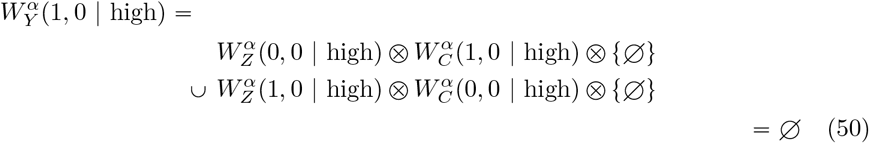

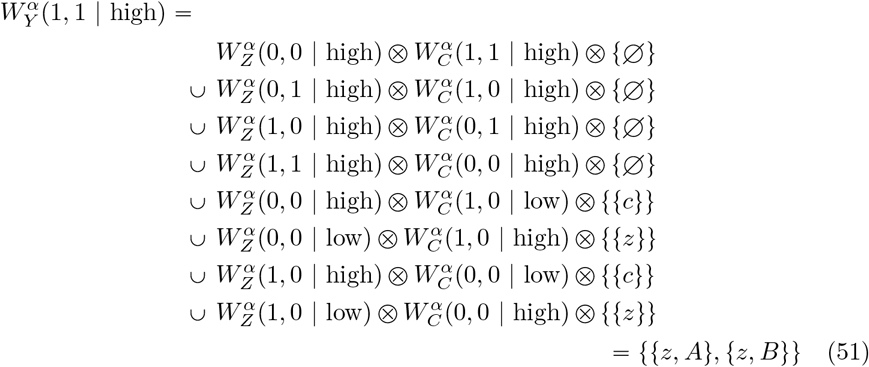

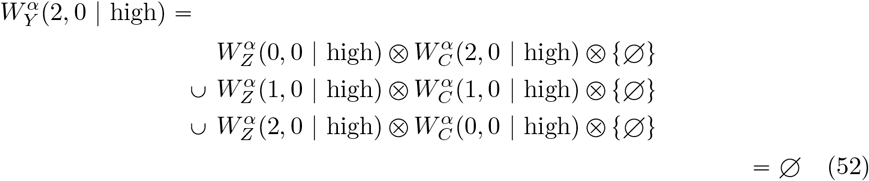

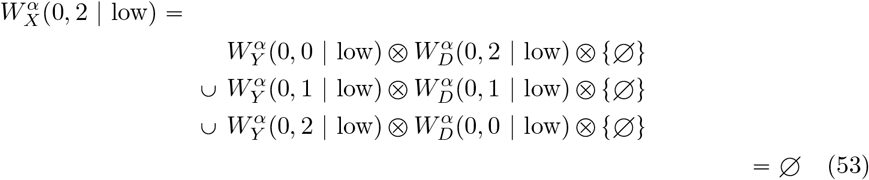

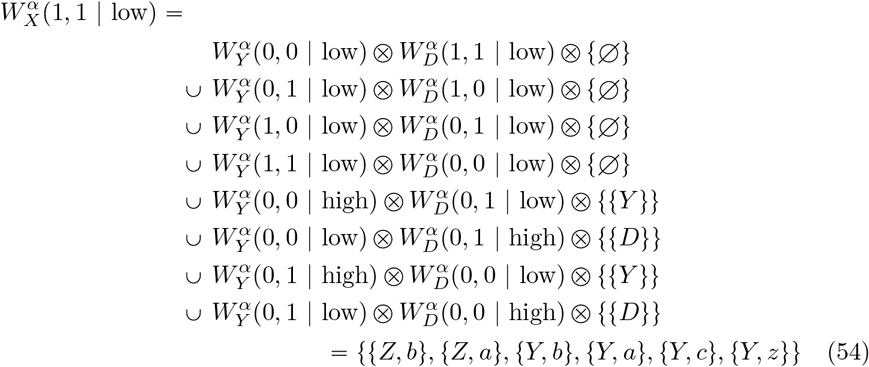

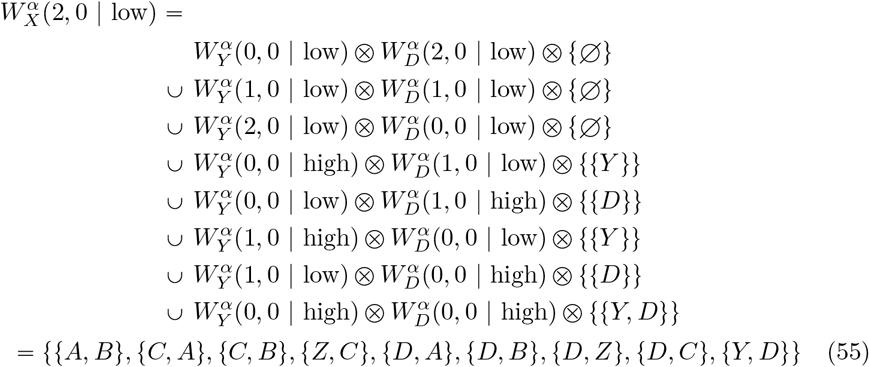

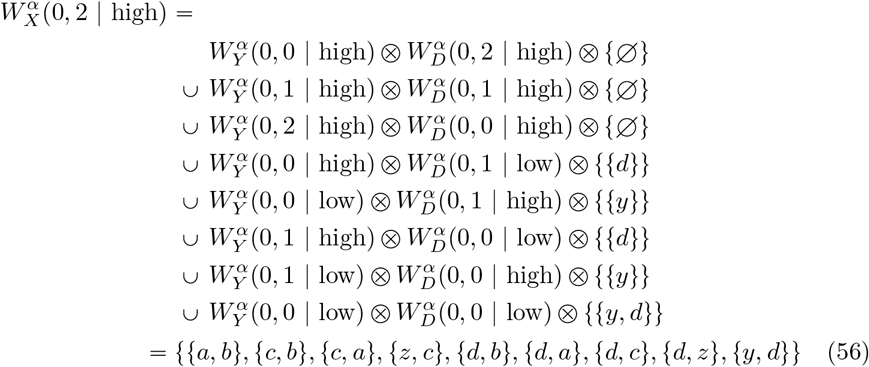

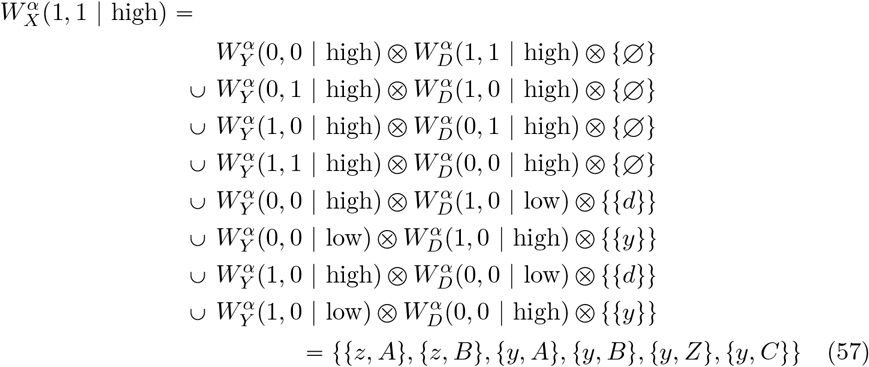

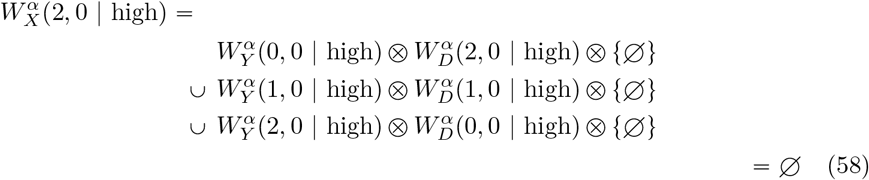

### Ways that one event in character *β* can happen in a tree

Given character *β* from Fig. 4b (Σ_*β*_ = {28, 30}), we detail how to compute all possible ways that one event, from *t*(Σ_*β*_) = {28→30, 30→28} can happen on tree *T* of Fig. 4a, given either state 28 or 30 at the root (*X*) of the tree *T*, i.e., 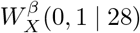 (Eq. 71), 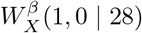 (Eq. 72), 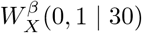 (Eq. 73) and 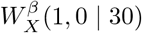 (Eq. 74). The base case(s) for the leaf nodes *A, B, C* and *D* of tree *T* is set according to Eqs. 12 as follows

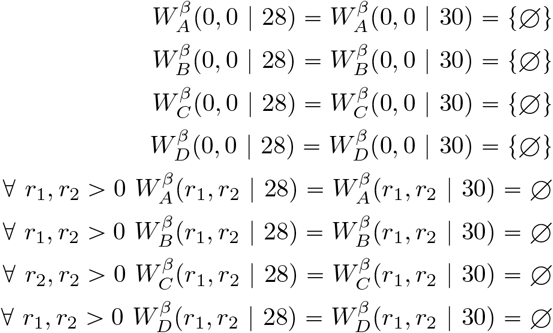

The recursive step(s) for each internal node *Z, Y* and finally *X* of tree *T* is then computed according to Eq. 13 as follows

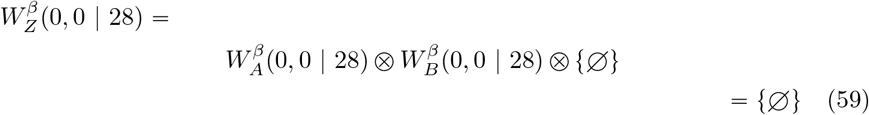

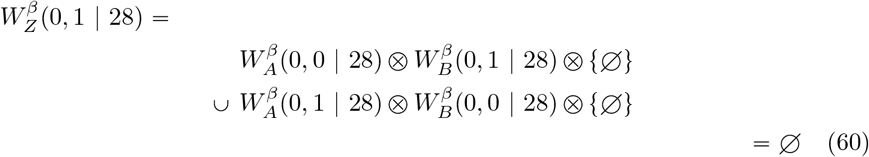

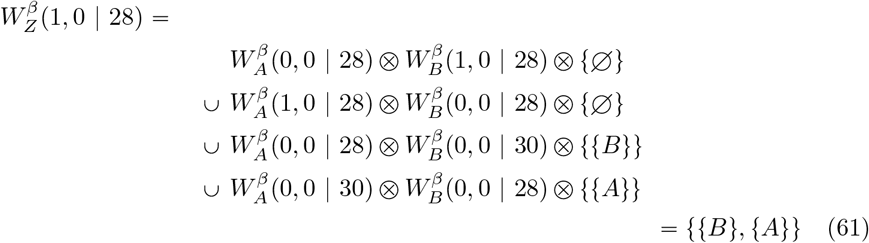

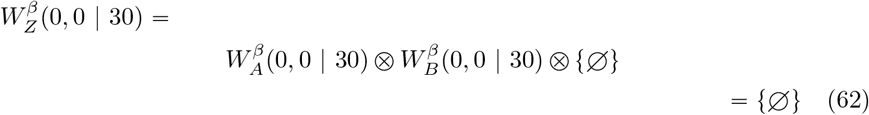

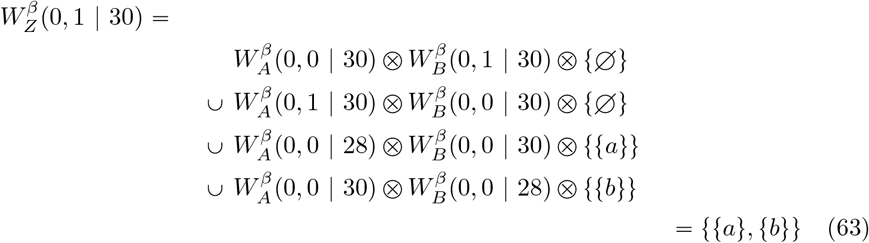

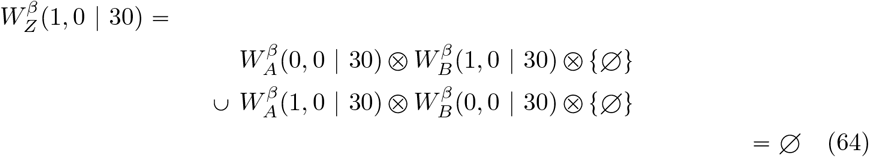

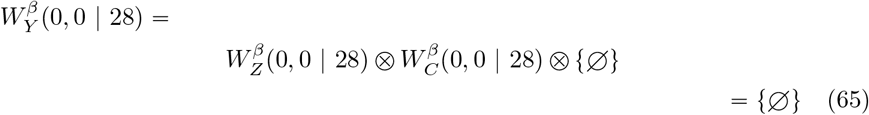

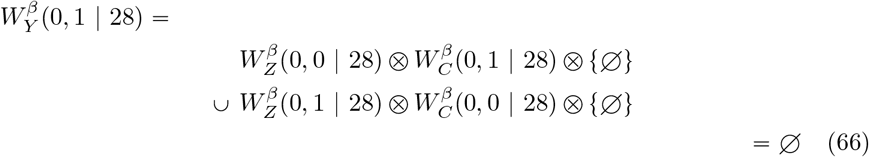

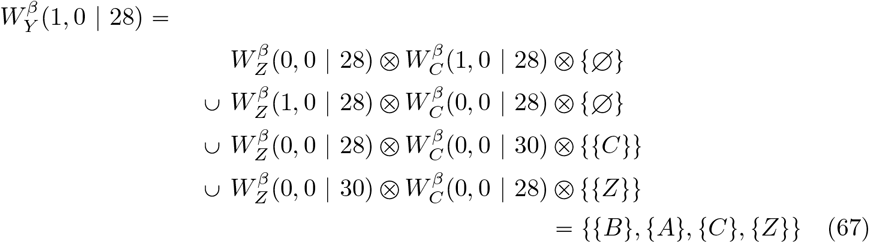

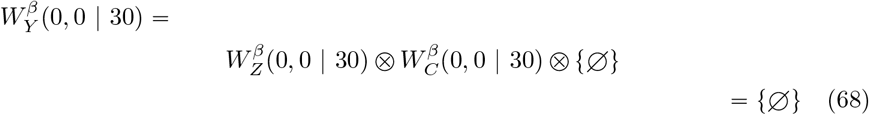

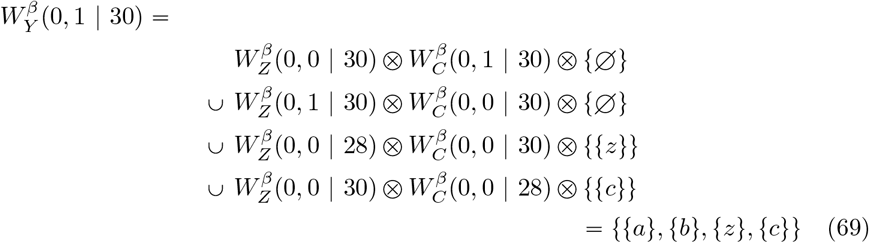

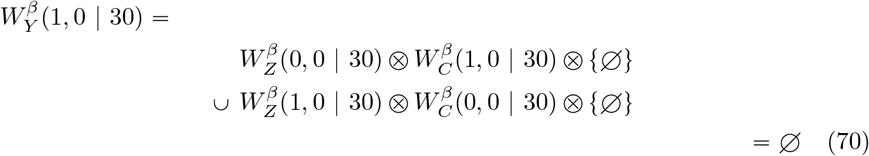

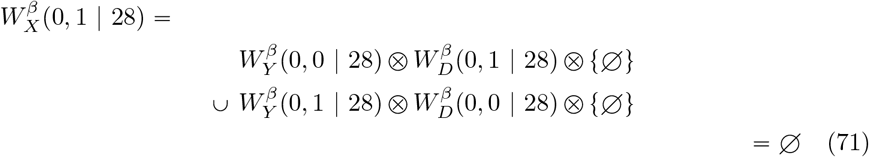

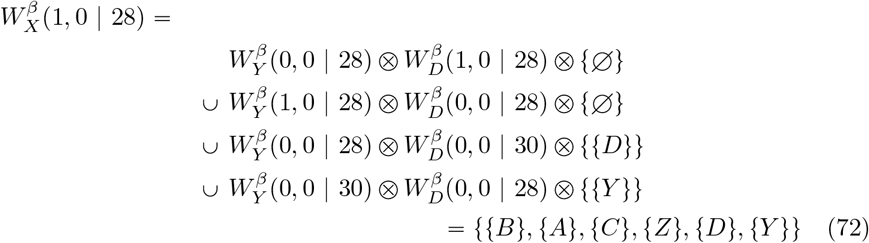

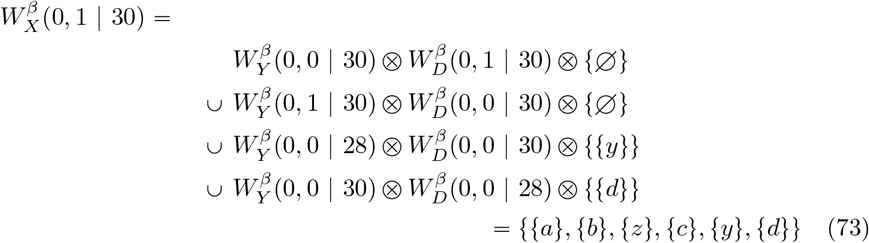

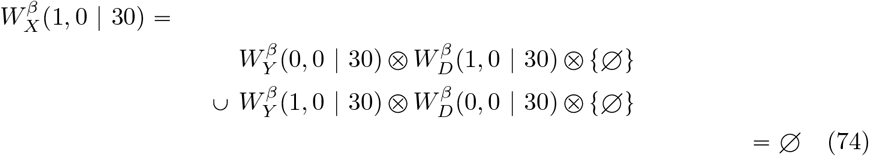

## Footnotes

1 A *large parsimony* is then a phylogeny which results in a small parsimony with the fewest changes (and this resulting small parsimony)

2 This is a generalization of the computation of *W* from [29], to sets of ways a character on any number of states can happen in a general (multifurcating) tree, in fact.

## Notes

https://github.com/murraypatterson/parcours

### Competing Interest Statement

The authors have declared no competing interest.

### Summary of Updates

1. changed way correlation was computed 2. different dataset 3. added way to perform significant correlation

